# The hippocampus encodes delay and value information during delay-discounting decision making

**DOI:** 10.1101/495598

**Authors:** Akira Masuda, Chie Sano, Qi Zhang, Hiromichi Goto, Thomas J. McHugh, Shigeyoshi Fujisawa, Shigeyoshi Itohara

## Abstract

The hippocampus, a region critical for memory and spatial navigation, has been implicated in delay discounting, the decline in subjective reward value when a delay is imposed. However, how delay discounting is encoded in the hippocampus is poorly understood. Here we recorded from the hippocampal CA1 region of mice performing a delay-discounting decision-making task, where delay lengths and reward amounts were changed across sessions, and identified subpopulations of neurons in CA1 which increased or decreased their firing rate during long delays. The activity of both delay-active and -suppressive cells reflected delay length, reward amount, and arm position, however manipulating reward amount differentially impacted the two populations, suggesting distinct roles in the valuation process. Further, genetic deletion of NMDA receptor in hippocampal pyramidal cells impaired delay-discount behavior and diminished delay-dependent activity in CA1. Our results suggest that distinct subclasses of hippocampal neurons concertedly support delay-discounting decision in a manner dependent on NMDA receptor function.

## Introduction

Animals faced with multiple options optimize their decisions through a complex cost-benefit valuation. The introduction of a time delay decreases preference for the delayed option (delay discounting) (Ainslie, 1975, 1992), with the discount rate varying on an individual basis. People who are considered patient exhibit lower discount rates, while impatient (or impulsive) people exhibit higher discount rates. Further, higher discount rates have been shown to be related to various neuropsychological disorders (Bickel and Marsch, 2001; Chesson et al., 2006; Epstein et al., 2008; Luman et al., 2010; Odum et al., 2000; Weller et al., 2008). While lesion studies have revealed a critical role for the hippocampus in delay discounting (Figner et al., 2010; Kalivas and Volkow, 2005; Peters and Büchel, 2011; Cheung et al., 2005; Mariano et al., 2009; McHugh et al., 2008), how this is reflected in hippocampal activity remains poorly understood.

Decades of study point to a critical role for the hippocampus in episodic memory (Scoville and Milner, 1957; Squire, 1992) and spatial navigation (Burgess et al., 2002; Ekstrom et al., 2003). While much of the rodent hippocampal physiology literature has focused on the spatial code present in hippocampal place cell activity (Jung and McNaughton, 1993; O’Keefe and Dostrovsky, 1971; Wilson and McNaughton, 1993), subsequent work has demonstrated that the circuit is capable of encoding a variety of features beyond the animal’s current position, including past and future trajectories (Ambrose et al., 2016; Foster and Wilson, 2006; Johnson and Redish, 2007; Johnson et al., 2007; Pfeiffer and Foster, 2013; Skaggs and McNaughton, 1996), the location of other animals or objects (Danjo et al., 2018; Omer et al., 2018), internal time (MacDonald et al., 2011, 2013; Manns et al., 2007; Pastalkova et al., 2008), and various physical scales (Aronov et al., 2017; Terada et al., 2017). Studies combining imaging and recording with optogenetic manipulation and identification suggest that subsets of CA1 neurons can encode distinct features of a task (Cembrowski et al., 2016; Danielson et al., 2016). Consistent with these data Gauthier and Tank recently identified a unique population of neurons which are active at reward sites, serving as “reward cells” (Gauthier and Tank, 2018). Although the reward-based activity has been well-investigated in relation to spatial context (Hölscher et al., 2003; Murty and Adcock, 2014; Ólafsdóttir et al., 2015; Peters and Büchel, 2010; Singer and Frank, 2009), the impact of the introduction of delay, the core computation for delay-based decision making, has not been examined. Thus, a central question is whether the activity of a distinct population of neurons in hippocampus is dependent on both reward-size and delay length.

Here we designed experiments to identify and characterize hippocampal neurons engaged during the delay of a delay-discounting task and to probe their sensitivity to changes in delay length and reward amount. We recorded singleunit activity in CA1 of mice performing a delay-discounting version of the T-maze task (Zhang et al., 2018) and assessed changes in neural activity related to delay length, arm position, and reward size. We found two populations of CA1 neurons which demonstrated increased or decreased delay-period activity. The activity of these both populations reflected delay length, reward size, and/or arm positions, however manipulation of reward size resulted in opposite responses between them, suggesting distinct roles in the overall valuation process.

## Results

### Behavioral profiling

We conducted a delay-based decision-making task in mice using the T-maze, where mice chose between right or left goal arms, with each arm containing a small reward or large reward, and with or without delay, respectively (Figure 1A). In total, we employed four behavioral conditions, with each session consisting of about 10 trials within 30 min, with a 20 s inter-trial interval at the start zone.

**Figure 1.**
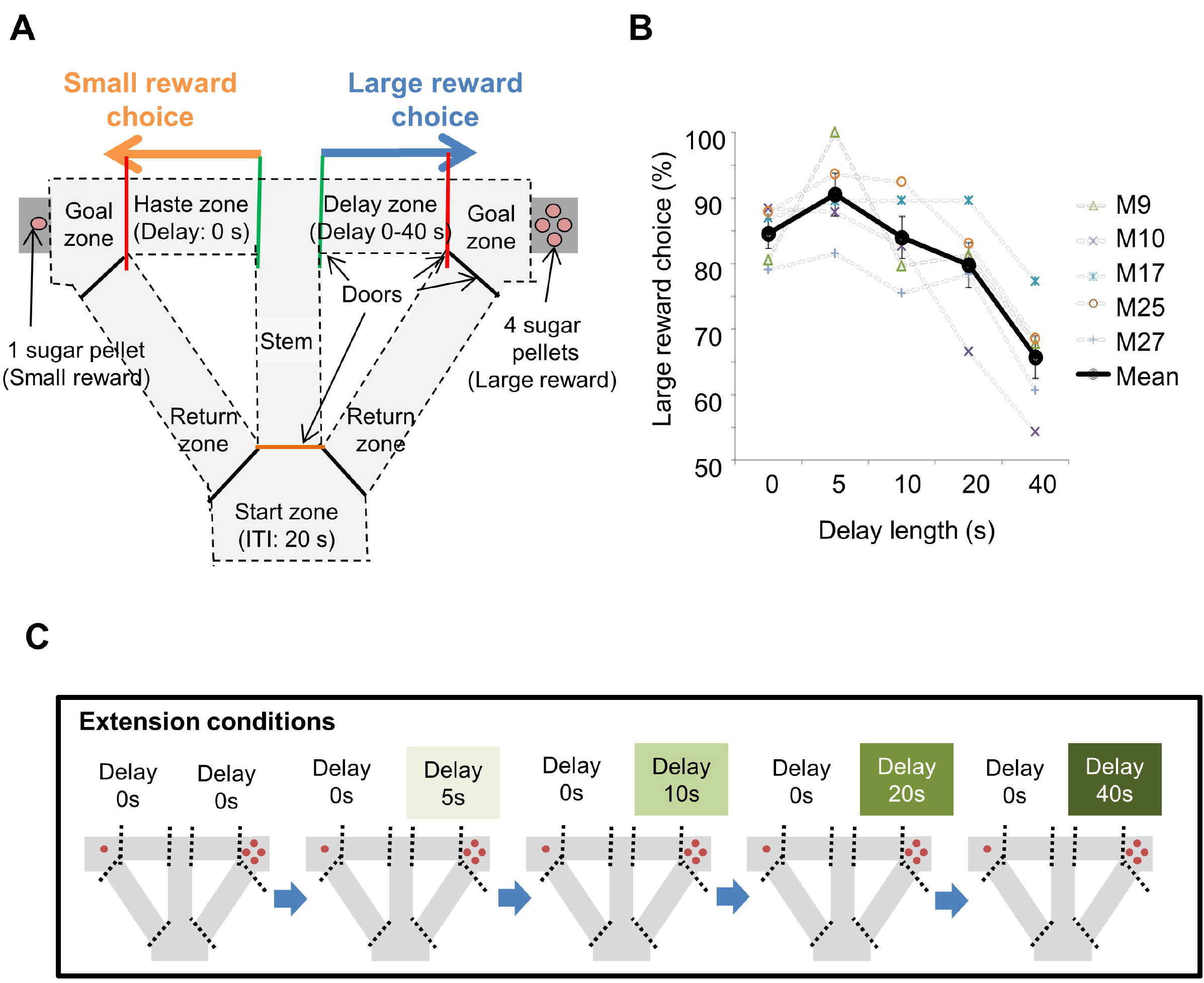
Task design of the delay-based decision making in the T-maze. (A) Schematic diagram for the experimental setup. Mice can choose the right or left arms assigned into the small reward without delay or large reward with delay, respectively. (B) Percentages of large-reward choices as a function of delay length. Error bars indicate the standard error of the mean (SEM). (C) Flow of the extension conditions. The delay lengths were extended sequentially. Red circles indicate the number of sugar pellets.

To examine the impact of delay on decision making, we changed the delay length sequentially. Once mice showed a preference for the large reward arm (> 80%) we increased the length of delay in a stepwise fashion (e.g., 0, 5, 10, 20, and 40 s; Figure 1C). With the inclusion of a delay, preference for the large reward arm decreased as a function of delay length (Figure 1B).

### Delay-dependent neuronal activity in the CA1

We recorded extracellular single units and local-field potentials (LFPs) from the CA1 region in total of 28 mice during delay-discount behavior and classified cells as putative excitatory neurons or inhibitory neurons based on characteristics of their extracellular waveform (Figure S2; see Materials & Methods). We first analyzed LFP signals in the CA1 region during delay periods. Consistent with the active movement of the mice, sharp-wave/ripples (SWRs) were rarely observed (*Z* = 1.97, *P* = 0.04, for start and stem zones vs delay zone, *Z* = 1.97, *P* = 0.03, for delay zone vs goal zone, Mann-Whitney *U* Test, Figure 2A and 2B) and the LFP was dominated by theta-range (7-11 Hz) activity (Figure 2C and 2D), suggesting the circuit remained engaged during this phase. We then examined the activity of excitatory neurons (Table S1) during specific task events, the exit from the start zone (start), the entrance/exit of the delay zone (delay), and the entrance to the goal zone (goal). In CA1 a subset of neurons exhibited delay-related activity, with firing rate rising during longer delays (> 20 s) (Figure 3A), while a distinct subset fired only under short delay conditions (Figure 3B).

**Figure 2.**
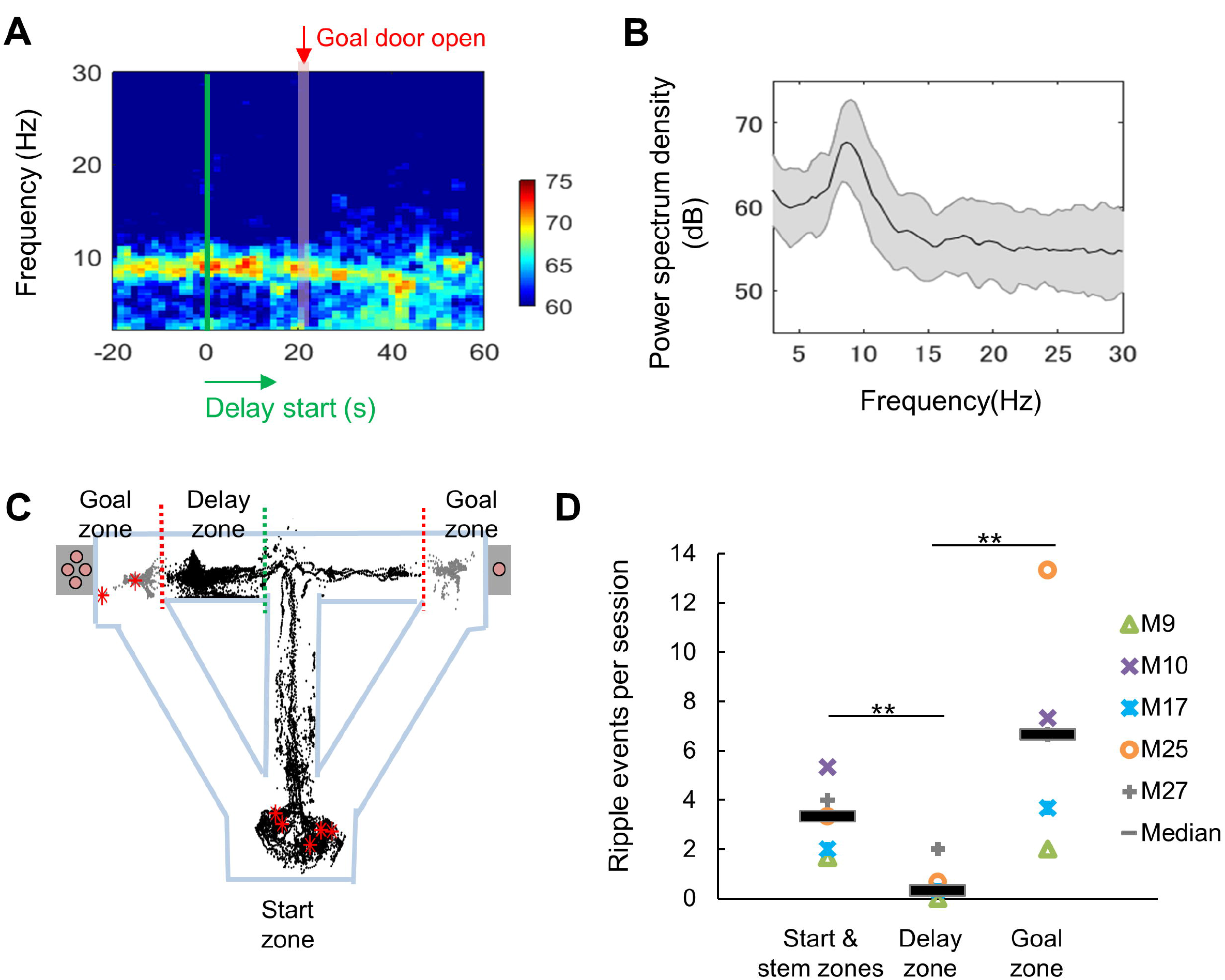
LFP signals during the long delay was characterized by strong theta power and lack of SWRs. (A) Sharp-wave/ripple events (SWRs) rarely occurred during the delay in the task (data from one session of delay 20 extension conditions). Red asterisks indicate the locations of SWRs. Black and gray dots indicate the path of animal movements before (black) and after arriving at the goal (gray), respectively. (B) SWRs per session in specific experimental zones. Total number of SWRs for each zone was counted and color-coded according to individual animals (average number of events acquired from 3 sessions of delay 20 extension conditions). *: *P* < 0.05, Mann-Whitney’s *U*-test. (C) Spectrogram of the hippocampal CA1 region at the peri-delay period (averaged from 3 mice, total 6 sessions of delay 20 extension conditions) Green line: delay-onset; red line: estimated goal-onset. (D) Power spectrum density during 2 s at the beginning of delay. Shaded area indicates ± SD.

**Figure 3.**
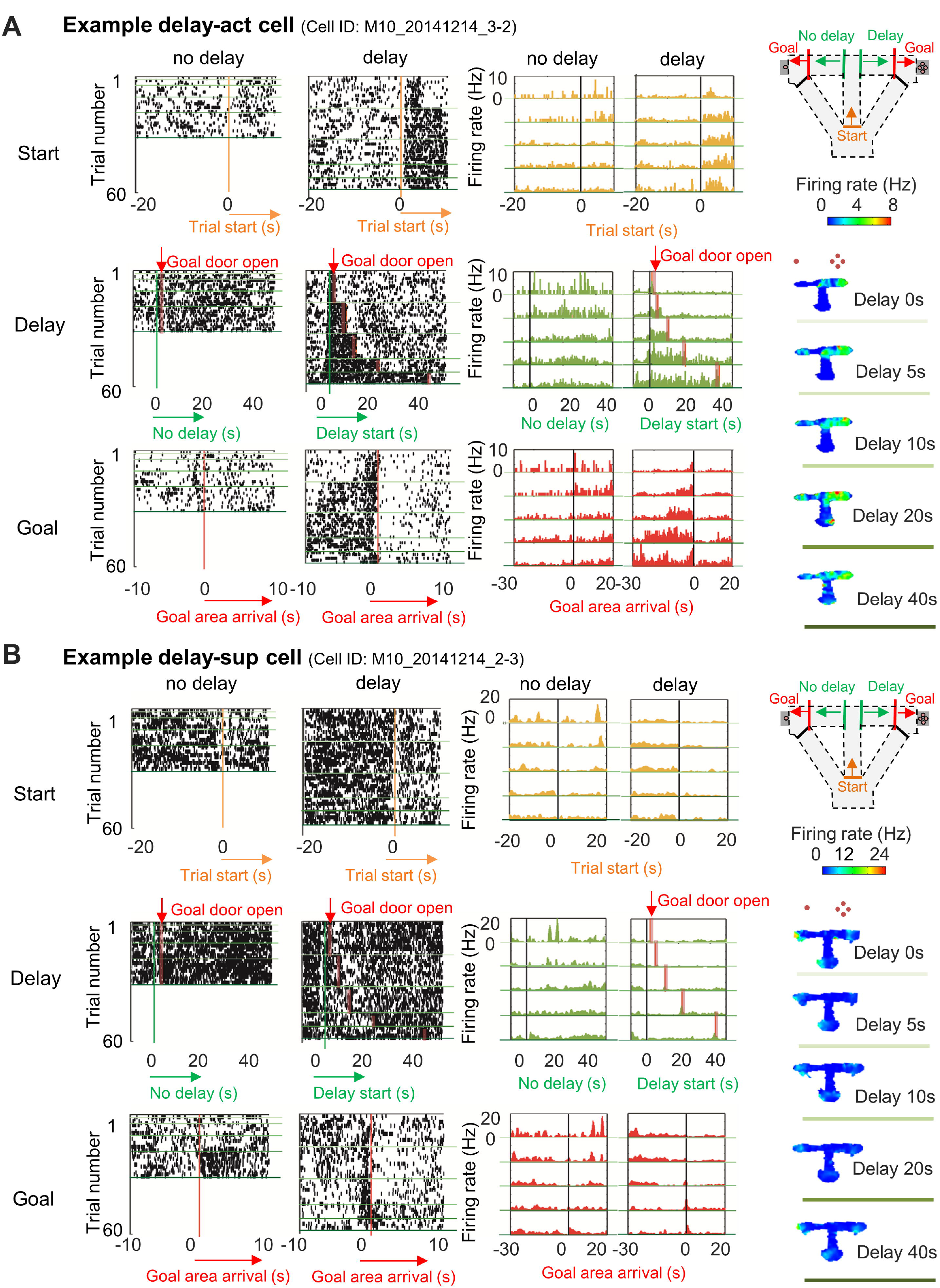
Increased or decreased neuronal activity of CA1 cells during delay. (A) An example of CA1 delay-active cells, which showed an increment in the firing rate as a function of delay length. Left, Raster plots of the firing activity of the cells aligned with start-onset (Top), delay-onset (middle) and goal-onset (bottom). Orange lines indicate start-onset. Green lines indicate delay-onset. Pale red lines indicate expected delay-offset. Red lines indicate goal-onset. Center, PSTHs of the firing activity of the cells aligned with start-onset (Top), delay-onset (middle) and goal-onset (bottom). Right, Color-coded rate maps. The delayed arm was assigned to the right side with a large reward for this recording session. Red dots indicate the number of sugar pellets. (B) An example of CA1 delay-suppressive cells, which showed a decrement in the firing rate during delay. The delayed arm was assigned to the right side with a large reward for this recording session.

We next asked if neurons significantly altered their firing rate during long delays compared with other phases of the task (see Materials & Methods). We found that across all task conditions large numbers of neurons exhibited significant increases (CA1: 243/639 units: 38.0%) or decreases (CA1: 313/639: 48.9%) in their firing rates during the delay (Table S1). We termed these delay-active (delay-act) and delay-suppressive (delay-sup) neurons respectively (Figure 4A). Average firing rates of delay-act and delay-sup neurons are shown as a peri-stimulus time histogram (PSTH) aligned at the onset of the delay (Figure.4B). When we examined the activity of neurons sequentially recorded under all possible delays we found the lengthening the delay dynamically altered the mean firing rates, with a subset of units (3/58: 5.1 % for delay-act; 8/83: 9.6 % for delay-sup; Table S2) demonstrating a significant correlation between the firing rate and delay lengths (Figure 4C and 4D). Comparison between the firing rates for short delays (5 s) and those for long delays (20–40 s) revealed that some delay-act and delay-sup cells exhibited significant elevation or reduction of firing rates for specific delay lengths (Figure 4E). At the population level, peak firing times of both CA1 delay-act and -sup cells were highly distributed when entering the delay zone (Figure 5A). To assess the population activity of CA1 cells during the task, we examined the autocorrelation of the population vectors under long delay conditions (Figure 5B). The population activity of both delay-act and -sup cells was clearly segmented in three periods, start, delay and goal, and with differential patterns of sustained activity in each.

**Figure 4.**
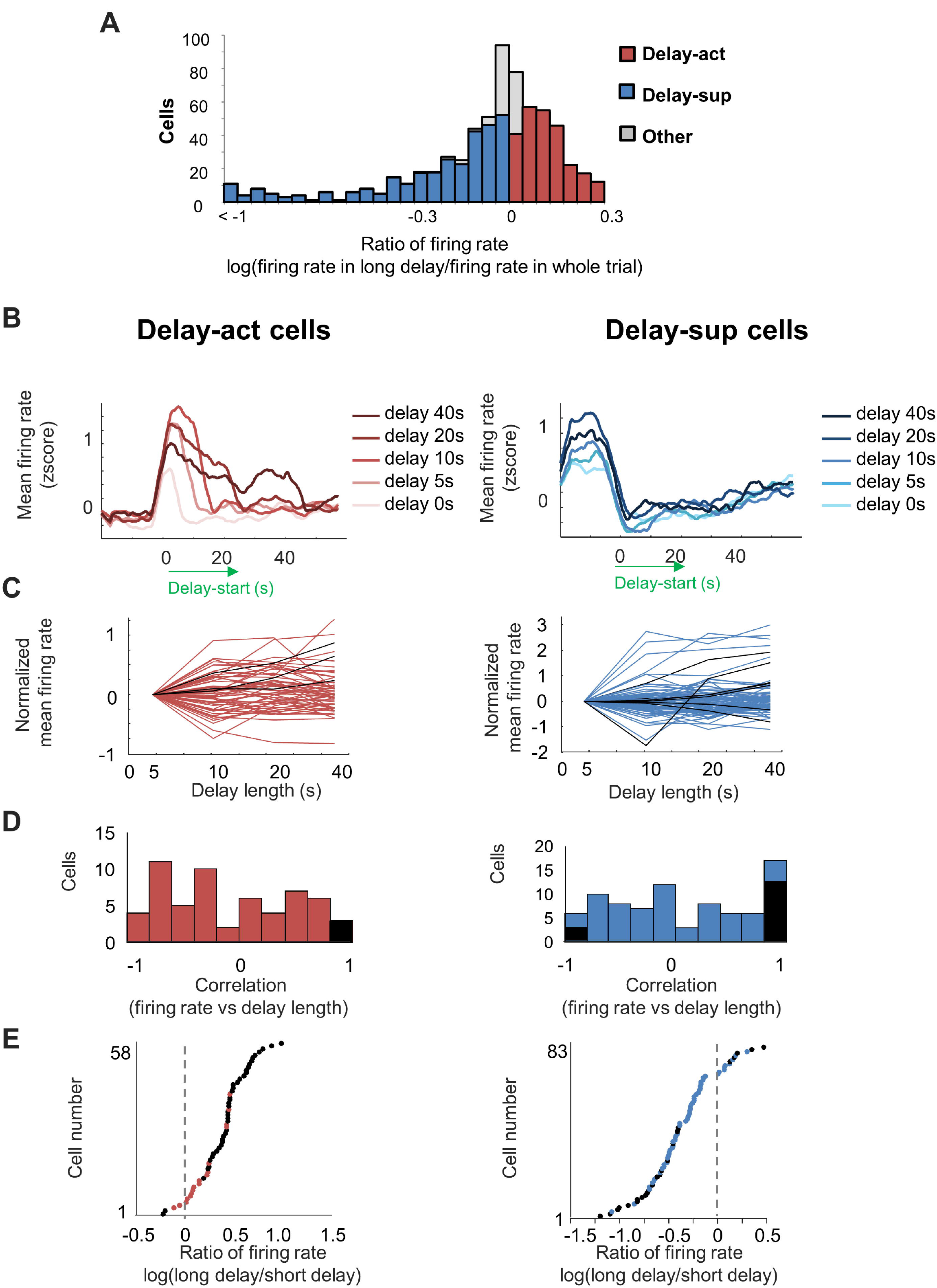
Delay-dependent firing patterns of CA1 delay-active and delay-suppressive cells. (A) The distribution of delay-active and delay-suppressive cells aligned with ratio of firing rate between in the long delay period and in whole trials. (B) Average firing patterns of all CA1 delay-active and delay-suppressive cells for different delay lengths (0, 5, 10, 20, and 40 s) (Extension conditions). Data were expressed as z-scored values. (C) Changes in firing rate as a function of delay length. Log-transformed ratios of firing rate (standardized by averaged the firing rate of delay 5 conditions) for CA1 delay-active and -suppressive cells are shown individually. Black lines indicate cells with statistically significant correlations with delay lengths (P < 0.05). (D) The distributions of correlation coefficients between the firing rate and delay length in CA1 delay-active and -suppressive cells. Black areas indicate cells with statistically significant correlations (P < 0.05). (E) The ratio of firing rate at long delay (20 or 40 s) to that of short delay (5 s) for all neurons (base-10 log-transformed). Each dot indicates an individual neuron. Black dots indicate neurons with a statistically significant difference in firing rate between short and long delay conditions (P < 0.001).

**Figure 5.**
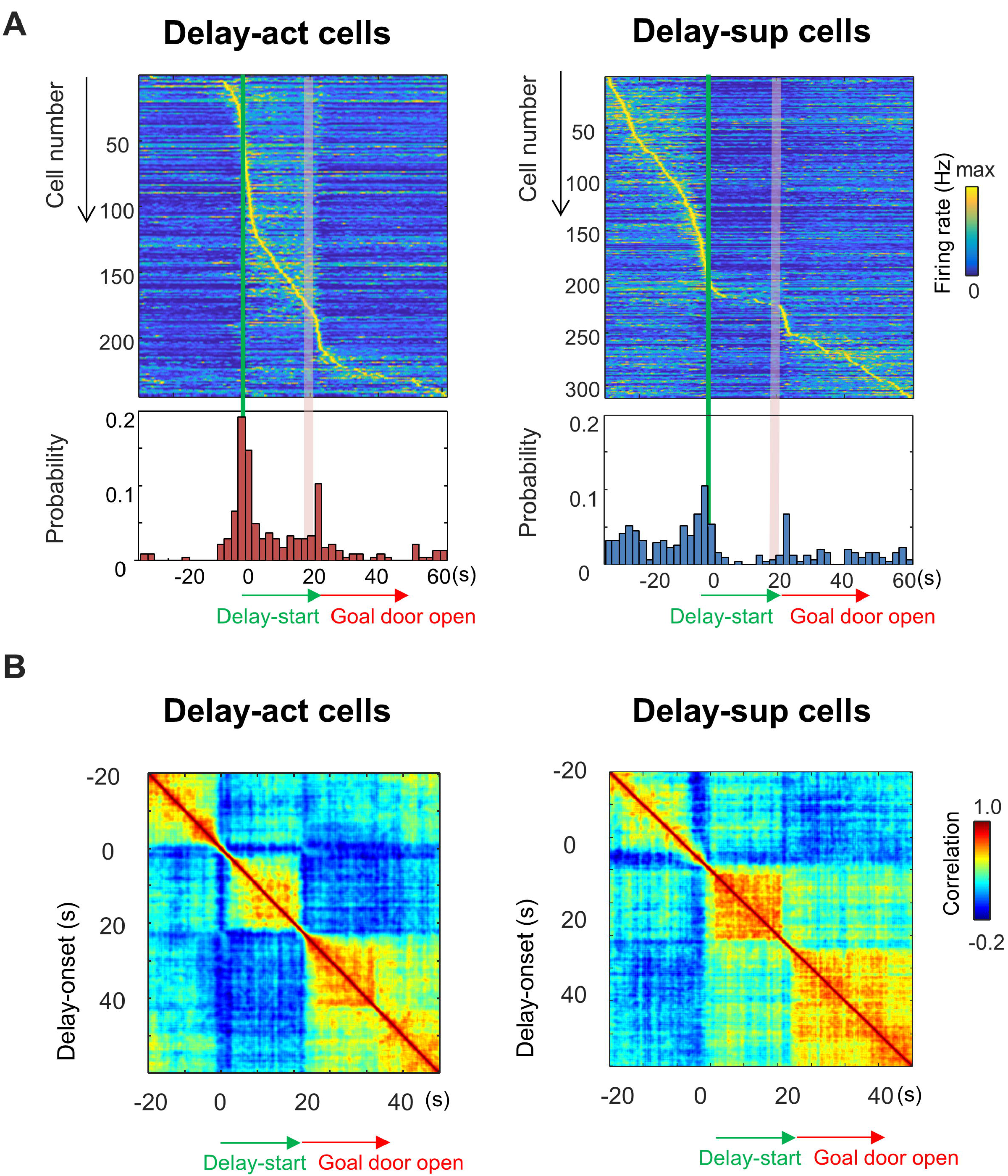
Temporally segmented population coding in CA1 delay-active and delay-suppressive cells. (A) Temporal patterns of firing rates in CA1 delay-active and -suppressive cells during delay. Top, Color-coded temporal firing patterns. Neurons were ordered by the time of their peak firing rates. Bottom, temporal distribution of peak firing rates of the neurons. Green lines indicate delay-onset. Pale red lines indicate expected delay-offset. (B) Correlation matrix of population vectors as a function of time for CA1 delay-active and delay-suppressive cells.

At the beginning of the daily recording session, about 10 % (4/33) of delay-act cells initially fired after the animal reached the goal in the 0 s delay condition, then shifted their firing to the delay period once a delay was introduced (Figure S3). Interestingly, subsequent elimination of the delay did not result in return to goal related activity (Figure S3A). We then compared the firing of CA1 delay-act cells under pre- and post-long-delay-conditions (Figure S3B). About 10% of neurons shifted positively or negatively (Figure S3C), indicating that the onset of firing in the CA1 neurons was influenced by the experience of waiting or learning of the delay.

### Place-specific delay information is encoded in majority of CA1 neurons

Given the robust place code present in the hippocampus we next asked if CA1 delay-act neurons were spatially-selective. To this end, we switched the location of the delay and no-delay arms (“Switched conditions”) or replicated the delay on the other side (“Both-side conditions”) (Figure 6A), with corresponding changes in reward size. Under both conditions the mice changed their behavior within several trials, with the preference for the large reward arm reaching about 70%. We then evaluated side-selectivity of the delay activity, adding location as a variable under 3-way ANOVA (side, delay-length, and timing; see Materials and Methods) during switched and both-side trials (Figure 6A). Representative side-selective and -unselective excitatory neurons in the CA1 are shown in Figure 5C. The percentage of side-selective neurons was high in CA1 delay-act (114/155: 73.5%) and -sup (124/191: 64.9%) neurons (Figure 6B and Table S2 and S3), however more than a quarter of the neurons of both groups encode delay independent of location.

**Figure 6.**
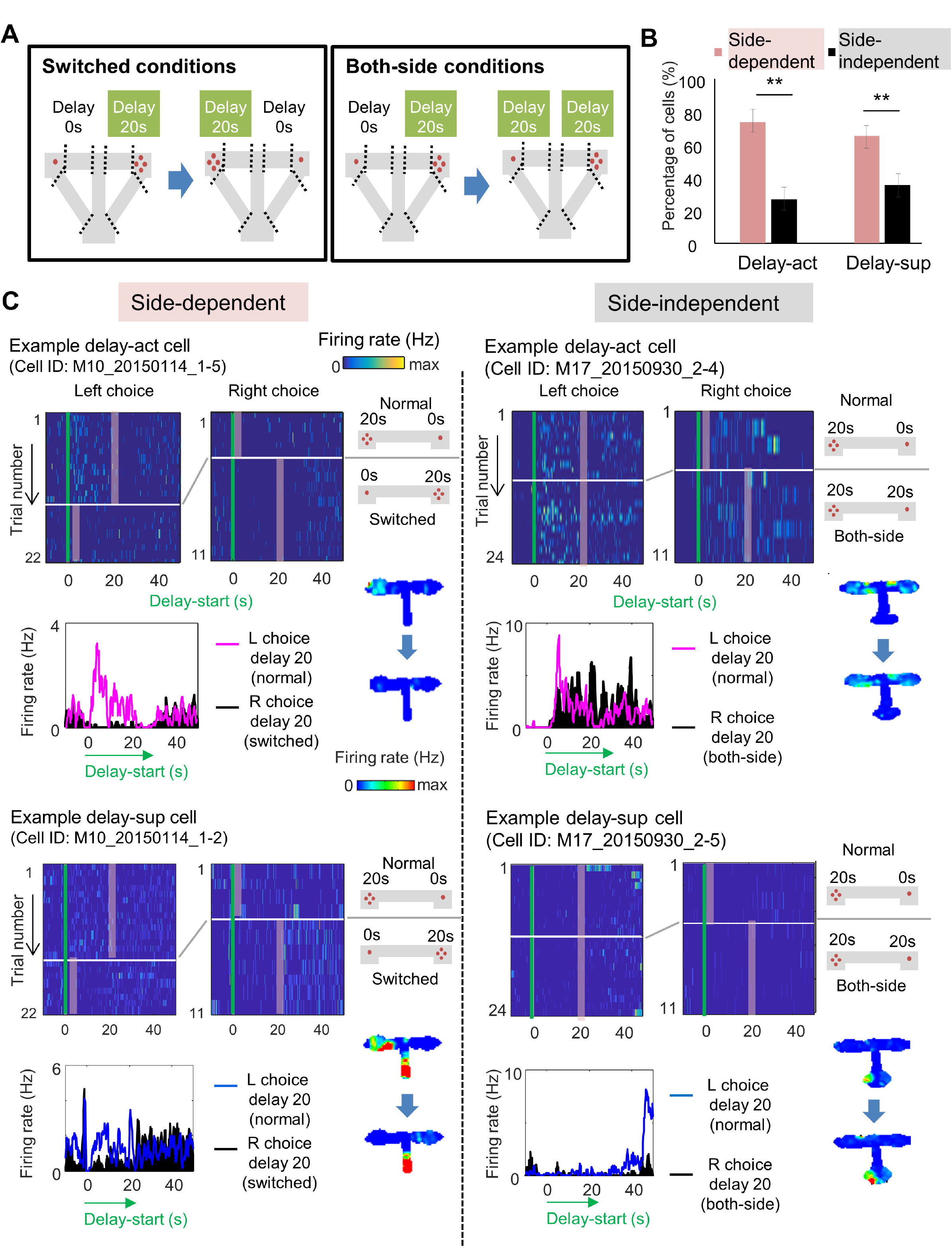
Spatial-selective delay coding in CA neurons. (A) Experimental conditions to investigate the location selectivity in delay-active neurons. The location of the delay zone was switched to the other side (switched conditions) or doubled to the both side (both-side conditions). (B) Percentage of place-dependent and-independent CA1 delay-active and delay-suppressive neurons. Error bars indicate 95% Clopper-Pearson’s confidence intervals. **: *P* < 0.01, Mann-Whitney’s *U*-test. (C) Example CA1 delay-active and delay-suppressive cells. Side-dependent and -independent neurons are shown as left and right rows, respectively. Top left, Colored raster plots expressing relative firing rates. Green lines indicate delay-onset. Pale red lines indicate expected delay-offset. Top right, Information of conditions corresponded to the raster plots on the left. Red dots indicate the number of sugar pellets. Bottom left, Peri-event time histograms showing the averaged firing rates. Magenta lines indicate the firing rate of the left choice with a 20-s delay. Black filled histograms indicate the firing rate of the right choice with a 20-s delay. Bottom right, Color-coded rate maps for the two conditions (normal delay and switch or both-side conditions).

### CA1 delay-active and delay-suppressive cells oppositely react to reward size manipulation

We further asked how subjective value influenced the activity of the delay-act and delay-sup neurons in CA1. To examine whether the firing patterns were correlated with reward size increment and decrement, we first devalued the reward size for the delayed option (devalued conditions), and subsequently restored the larger reward (revalued conditions, Figure 7A) to avoid order-dependent confounds arising from decreased hunger or motivation of the animals in later trials. The majority of delay-act cells decreased their firing rate in response to reward devaluation, whereas delay-sup neurons had the opposite response, increasing their activity (Figure 7B and 7C). As a result, the log ratio of the firing rates (large reward/small reward) under both devaluated and revalued conditions was significantly different between delay-act and delay-sup cells (*Z* = 3.2, *P* = 0.001 for devaluation; *Z* = 5.9, *P* = 0.03 for revaluation, Mann-Whitney’s *U*-test). In total, the ratio was negatively skewed in the delay-act cells (*F* = -3.0, *P* = 0.004, one-sample *t*-test) but positively skewed in delay-sup cells (*F* = 2.6, *P* = 0.01, one-sample *t*-test). These results suggest that firing during the delay reflected positive and negative outcomes in different subpopulation of CA1 neurons.

**Figure 7.**
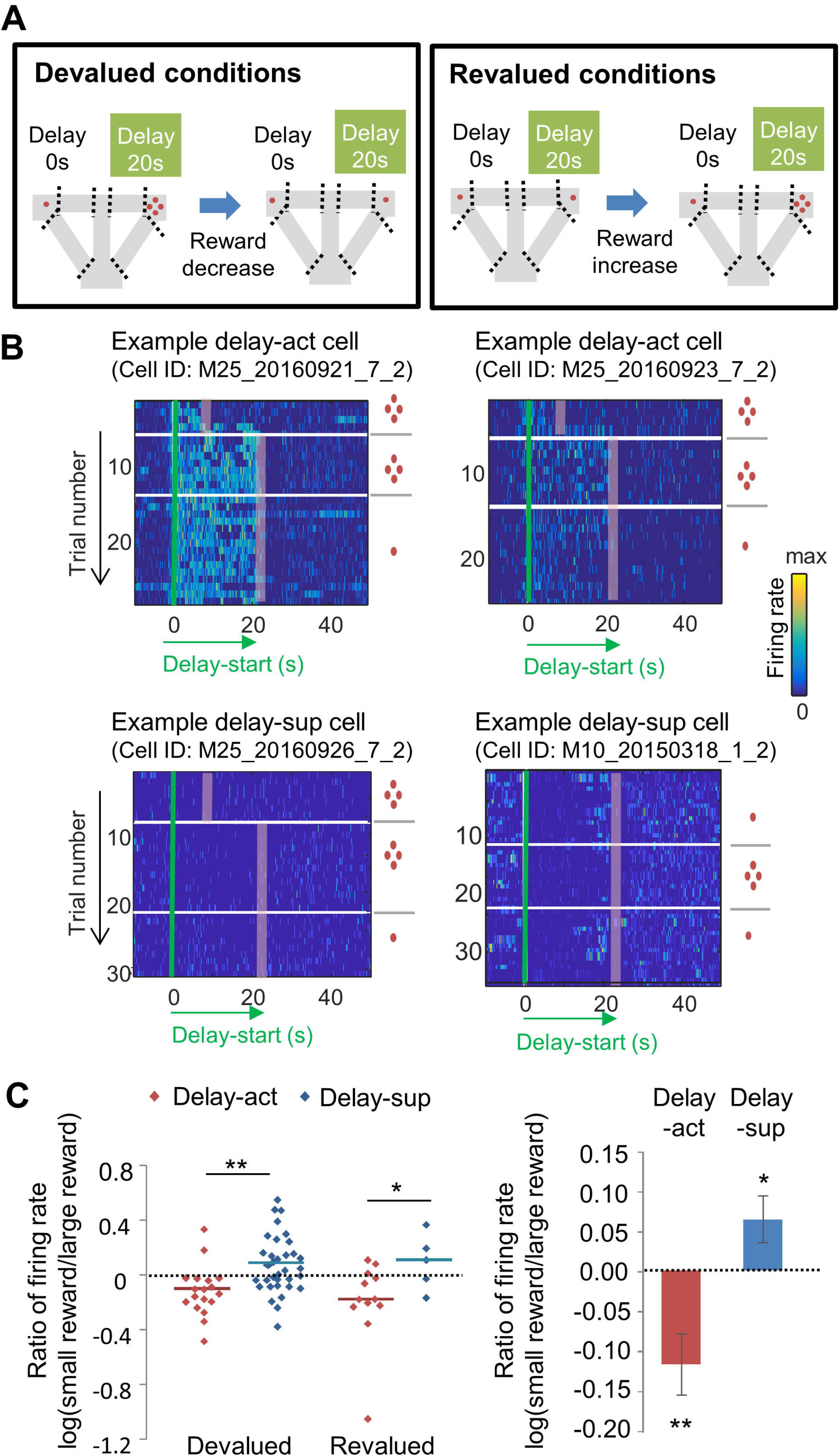
The firing of CA1.delay-active and delay-suppressive cells distinctly changed by reward size manipulations. (A) Left, Experimental conditions of the devalued conditions. The reward size was changed from 4 to 1 (or 0) pellet. Right, Experimental conditions of the revalued conditions. The reward size was changed from 1 (or 0) to 4 pellets. (B) Example CA1 delay-active (top) and delay-suppressive cells (bottom) fired during delay in devalued conditions. Green lines indicate delay-onset. Red lines indicate expected delay-offset. Red dots indicate the number of sugar pellets. Ratio of firing rates of delay-active and -suppressive cells in devalued and revalued conditions. Dots indicate individual data of delay-active cells (red) and delay-suppressive cells (blue). Central bars indicate the medians. *: *P* < 0.05; **: *P* < 0.01, Mann-Whitney’s *U*-test. (C) Ratio of firing rates of delay-active and -suppressive cells in mixed population. Error bars indicate SEM. *: *P* < 0.05; **: *P* < 0.01, One-sample t-test.

We next focused on the relationship between the behavioral shift over devaluation sessions and the firing patterns of delay-act cells. Across the session animals avoided the delayed option. To eliminate the preference for the delayed options, we designed an “unequal conditions” (long delay + no pellet vs long delay + 4 pellets, with the latter being the better option). Animals then quickly reduced their preference to the delayed option with no reward. To record the activity for the less-preferred or adverse choice, we forced mice to choose the less-preferred option with an obstacle set at the entrance of the opposite arm. When faced with an unrewarded delayed option CA1 neurons indicating choice preference were silent (Figure S4). These results suggest that the firing of delay-act neurons in the CA1 region represent the animal’s subjective value of the chosen options.

### NMDAR deficiency in hippocampus disrupted delay-discounting and populational delay coding in CA1

Finally, we took advantage of a mutant mouse, the *CaMK2-Cre; NR1-flox/flox* mouse, lacking CA1 pyramidal cell NMDA receptors (NMDARs) (CA1-NR1cKO mouse; McHugh et al., 1996; Tsien et al., 1996a), to assess the role of synaptic plasticity in task performance. Consistent with previous reports of hippocampal-dependent learning deficits in these mice (Bannerman et al., 2012; Rondi-Reig et al., 2001; Tsien et al., 1996b), they exhibited impaired delay discounting (Figure 8A), demonstrating a significant bias for the larger reward even when the delay was extended (F1 = 14.3, *P* < 0.001, genotype (CA1-NR1cKO vs NR1 f/f); F4 = 23.0, *P* < 0.001, delay length, F1,4 = 1.07, *P* = 0.37, two-way ANOVA, *P* = 0.61 on delay 0, *P* = 0.03 on delay 5, *P* = 0.04 on delay 10, *P* = 0.002 on delay 20, *P* = 0.005 on delay 40, multiple comparisons on each delay length).

**Figure 8.**
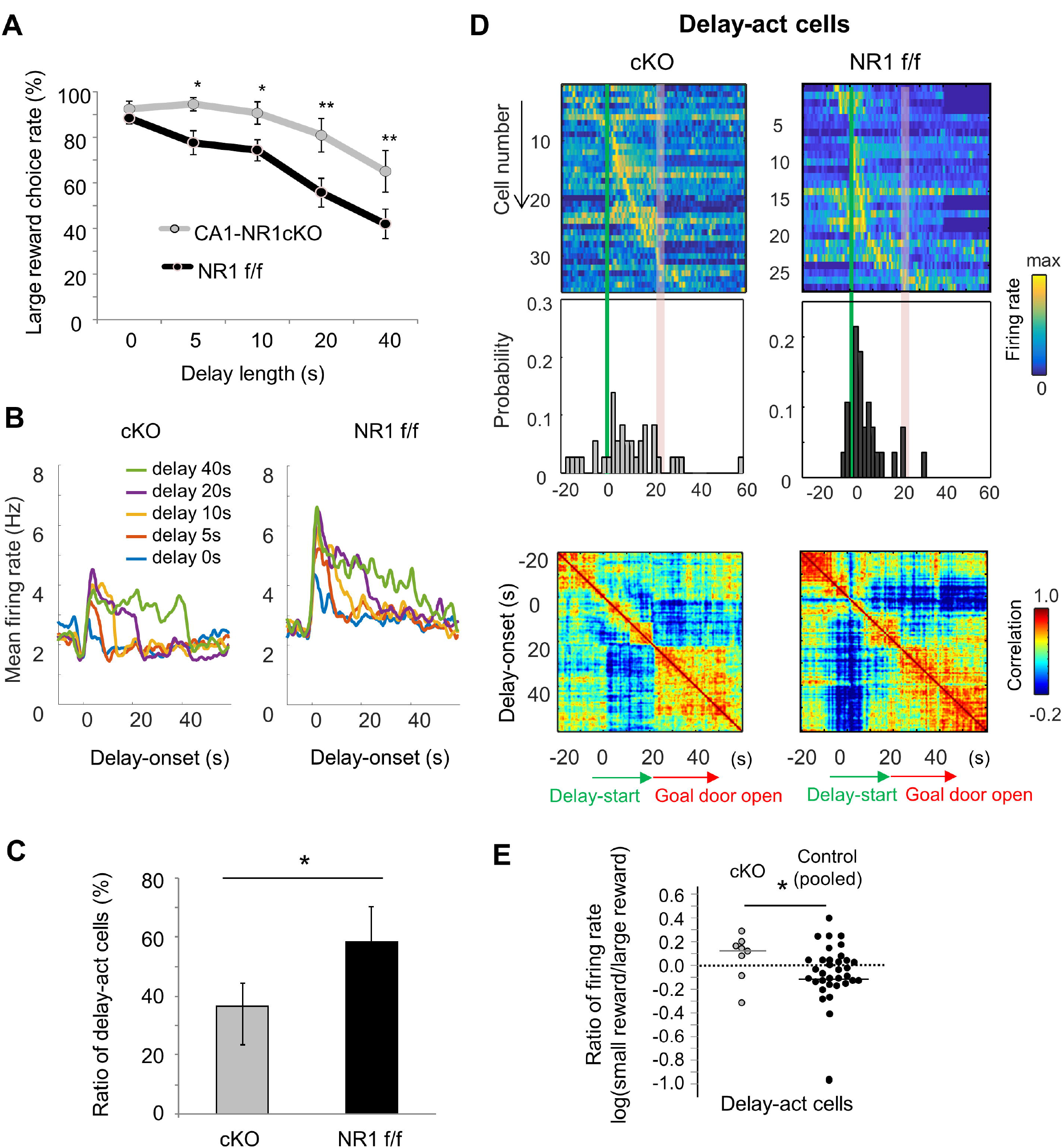
NMDAR-dependent mechanism for delay-discounting. (A) Impaired delay-discounting in CA1 -NR1 cKO mice. *: *P* < 0.05; **: *P* < 0.01, post-hoc Scheffe’s test. Error bars indicate SEM. (B) NMDAR deficiency disrupted the delay tuning in the CA1 activity. Top, Average firing patterns of the CA1 delay-active cells from cKO and control mice for different delay lengths (0, 5, 10, 20, and 40 s). Bottom, Abnormal delay-active and -suppressive cell proportion in cKO mice. Ratio of delay-act cells to delay-sup cells for cKO and control mice. Error bars indicate 95% Clopper-Pearson’s confidence intervals. *: *P* < 0.05, Mann-Whitney U test. (C) NMDAR deficiency disrupted the populational activity in CA1. Top, color-coded temporal firing patterns of the CA1 delay-active cells in cKO and control mice. Neurons were ordered by the time of their peak firing rates. Middle, temporal distribution of neurons. Green lines indicate delay-onset. Red lines indicate expected delay-offset. Bottom, Correlation matrix of population vectors as a function of time for CA1 delay-act cells in cKO and control mice. NMDAR deficiency disrupted the negative skew in firing rate ratio of delay-active cells. Ratio of firing rates of delay-active cells in CA1 of cKO and WT mice. Dots indicate individual data of cKO (gray) and control (black). Central bar indicate the median.

We next recorded CA1 neuronal activity in the cKO (n = 3, 123 units) and control mice (n = 3, 69 units, Table S4) to look for physiological correlates of the behavioral change. Delay-act cells in the cKO mice showed non-specific activation during the delay period and delay zone (Figure 8B and Figure S5A). Hence, there was a lower and higher proportion of delay-act and delay-sup neurons in the cKO respectively and the ratio of delay-act/delay-sup was significantly lower in the cKO compared the control mice (Figure 8C, *P* = 0.02, Fisher’s Exact Test). Further, in contrast to the controls, the temporal distribution of all delay-act cells in the cKO was sparse and not specific to delay-onset. As a result, population vector analysis revealed that the activity was not segmented into three periods in the cKO mice (Figure 8D). In addition, the ratio of CA1 firing of cKO was significantly different than that observed in control mice and lacked the expected negatively skewed distribution (*Z* = 2.0, *P* = 0.04, Mann-Whitney’s U-Test, Figure 8E). These findings suggest that delay discount behavior and the underlying delay-related activity in CA1 pyramidal cells requires NMDAR-dependent mechanisms in the hippocampus.

## Discussion

We recorded CA1 neuronal activity in mice during delay-based decision making in an automated T-maze task while independently manipulating delay length and reward size across sessions. We observed distinct populations of neurons that increased or decreased their firing during the delay. Moreover, the firing rates of a subset of the delay-activated CA1 neurons decreased with both delay length increments and reward size declines. Notably, the activated and suppressed neurons showed distinct activity changes following reward size manipulations.

These results suggest that dissociable subpopulation of hippocampal neurons represent delay and reward information in opposing ways. These discoveries should help shape models of how the hippocampus support decision making.

Although the delay-modulated firings were diverse in CA1 neurons, their responsiveness to delay was precisely controlled. Peak firing rates of delay-act and delay-sup cells were distributed largely around the delay-onset. As a result, population vector analysis demonstrated segmented and sustained network activity during the delay in the CA1 region, suggesting a role in prospective coding of specific periodic events centered on the delay.

A significant fraction of both delay-act and delay-sup neurons we recorded also carried spatially tuned delayed information. Thus, the activity of most delay-act and delay-sup cells in dorsal CA1 does not appear to represent solely delay information, but rather, may represent integrated information of the chosen option, reflecting both location and delay. This result is consistent with the idea that hippocampal cells are coding not only within the space and time dimensions individually, but rather across them jointly (Eichenbaum, 2014; Howard and Eichenbaum, 2015; MacDonald et al., 2011).

Changing reward size modulated the firing rates of both the delay-act and delay-sup cells in CA1. It is widely known that the activity of CA1 neurons can depend on reward (Ambrose et al., 2016; Hölscher et al., 2003; Singer and Frank, 2009). Studies focusing on goal-directed behavior have demonstrated that some CA1 neurons fire when animals approach, wait for, or acquire rewards, but not when animals visit the same location in the absence of the reward (Eichenbaum et al., 1987; Fyhn et al., 2002; Hok et al., 2007; Kobayashi et al., 2003; Rolls and Xiang, 2005), indicating that certain subset of CA1 neurons are highly sensitive to reward expectation or motivation. However, in monkeys, omitting a reward activated some CA1 neurons (Watanabe and Niki, 1985). This is consistent with our results demonstrating that during the delay, dissociable subsets of CA1 neurons were positively or negatively correlated with reward size. More importantly, we found that different subsets of CA1 activity reacted in an opposite manner to “delay increments” or “reward declines” (Figure 6C). Accordingly, distinct subpopulation of CA1 neurons may encode the delay-reward integration and may support valuation process in the delay-based decision making.

The phenotype of “lowered delay discounting” caused by a loss of the NMDAR may be interpreted also as an abnormal repetition of an unpleasant choice, referred to as “compulsive behavior”. Systemic injection of the partial NMDAR agonist D-cycloserine reduces compulsive lever-pressing in a model of obsessive-compulsive disorder (OCD) in rats (Albelda et al., 2010). In addition, polymorphisms in a subunit of NMDAR have been considered as a risk factor of OCD (Arnold et al., 2004). The present study provides evidence that hippocampal NMDARs may be associated with compulsive disorders.

Our results suggest that the hippocampal NMDAR is required for delay discounting; however, a key question of what the critical roles of hippocampal NMDARs are during delay discounting remains to be addressed. It is widely believed that synaptic plasticity via NMDAR-dependent machinery contributes to association learning, and that in the hippocampus contributes to the formation of long-term, spatial memories (Martin et al., 2000). Studies using several lines of conditional knockout mice have pointed out that NMDAR in the hippocampus is involved in spatial learning (Tsien et al., 1996a), nonspatial learning (Huerta et al., 2000; Rondi-Reig et al., 2001), anxiety (Bannerman et al., 2004; Kjelstrup et al., 2002; McHugh et al., 2004; Richmond et al., 1999), time perception (Huerta et al., 2000), and decision making (Bannerman et al., 2012). In addition, physiological studies have demonstrated that the hippocampus lacking NMDAR exhibits less specific spatial representation in place cells (McHugh et al., 1996). We found that NMDAR deficiency disrupted the proportion of delay-act and -sup cells, and population coding for the delay. These findings suggest that the NMDAR in the hippocampus may be required for maintaining or developing time-coding. Further research is required in order to identify more specific mechanistic roles of the hippocampal NMDAR in delay-based decision making. Additionally, in contrast to previous studies showing rats with hippocampal lesions exhibit higher discount rates (Cheung et al., 2005; Mariano et al., 2009; McHugh et al., 2008), the NMDA KO mice demonstrated the opposite phenotype. Thus, there may be considerable differences between the effect of the lesions and NMDAR knockout in the hippocampus on the full network engaged during delay-based decision making.

In conclusion, our results show that CA1 neuronal activity during delay is segregated into two populations, delay active and delay suppressive neurons. Further, these groups demonstrate opposing responses to changes in motivational background. In addition, NMDAR-dependent plasticity mechanisms appear to be required for the formation of the firing patterns during delay and for the delay-discounting. These findings further clarify the role of the hippocampus in decision making, as well as in the control of impulsive/compulsive behaviors.

## Materials & Methods

### Animals

A total of 28 male C57B6/J mice were used for this study [Wildtype: n = 5, cKO: n = 11 (8 for behavioral study); control: n = 12 (9 for behavioral study)]. Mice lacking NMDAR in the hippocampus were generated by crossing the line gene-targeted for loxP-tagged *Nr1* (*Grin1*) alleles (*Nr1*^flox^, Tsien et al., 1996) and a transgenic line carrying *Camk2a* promoter-driven Cre recombinase (*Camk2a-Cre*, T29-1Stl,Tsien et al., 1996a). In this mutant, deletion of NR1 is delayed until about weeks after birth and is restricted to the CA1 pyramidal cells until about 2 months of age (Fukaya et al., 2003). Most of the behavioral analysis using the mutant was done until this age. Hence, it is unlikely that the behavioral impairment observed was the result of undetected developmental abnormalities. Physiological characterization, however, may have harbored a more widespread deletion of the NR1 gene.

### Delay-based decision-making task

Adult mice were trained in a delay-based decision-making task under an automated T-maze (O’HARA & Co., Tokyo, Japan) before electrophysiological recording. The maze was partitioned off into 6 areas (Start, Junction, Right-Goal, Right-Back, Left-Goal, and Left-Back) by 7 sliding doors (S-J, J-R, R-RG, RG-S, J-L, L-LG, and LG-S). The detailed protocol has been described previously (Kobayashi et al. 2013). In short, the mice had food restriction to approximately 80% of free-feeding weight, were habituated to the maze, and baited with scattered pellets (30 min/day) for 2 days. The large reward arm and the small reward arm were allocated to the right or left side arm randomly among mice. Four pellets were available in the large reward arm, while only 1 pellet was available in the small reward arm. Mice were allowed to roam freely and without delay to select either arm for 5-10 days for the initial training period until they preferred the large arm (> 80%). Then, all animals were trained in the Extension delay conditions for at least 5 days. At the first block of trails for each day, the large reward arm was associated without delay (0-s), and then at the later blocks it was associated with a 5-s, 10-s, 20-s, or 40-s delay. In the meantime, the small reward arm was always associated with no delay. Each block consisted of 10 trials or more (15 or 30 min). If the trial number was lower than 10, additional blocks were employed. Next, the mice, except cKO and control, were trained in the Switched and Both-side conditions. In the Switched condition, the side of the delayed-large arm was switched to the other side. In the Both-side condition, both two sides were set as delayed-small and delayed–large arms. The Switched conditions were performed initially and then Both-side conditions. In changing the conditions, 10 or more trials were continuously performed to develop a sustained reaction in the animals. Finally, the mice were trained in the Devalued and Revalued conditions. We devalued the reward size to investigate whether the firing rate reflected a positive or negative aspect in the delayed option. Initially we set a delay for a short time with the normal large reward, and then we changed the delay to be long without any change of the large reward, similar to other conditions. After these two continuous sessions, we changed the reward size from four to one pellet. As other control conditions, we also performed the opposite flow (long delay with one pellet first, long delay with four pellets next). During the time between blocks, mice were allowed to drink water.

### Histological identification of the localization of the recorded sites

Due to the small thickness of the silicon probe shanks, the tracks of shanks were hard to detect. Painting at the back of the shanks and/or the electrical lesion by a small current (5 mA for 5 s) was used to facilitate track identifications.

### Recording & spike sorting

Mice were anesthetized with isoflurane during surgery. Silicon probes or wire tetrodes were implanted in the hippocampal CA1 region (AP = –2.0 to –2.8 mm, ML = 1.2 to 2.0 mm, DV = 1.2 to 1.5). In all experiments, ground and reference screws were fixed in the skull atop the cerebellum. The silicon probes were attached to micromanipulators (nDrive, NeuroNexus, Michigan, USA), which enabled us to move their positions to the desired depth. Electrophysiological signals were acquired continuously at 20 kHz on a multi-channel recording system (KJE-1001, Ampliplex Ltd, Szeged, Hungary). The wide-band signal was down-sampled to 1.25 kHz and used as the LFP signals. We detected SWRs (their timing, power, and durations) from filtered signal (120-230 Hz) that corresponded to larger than 3 SD of log-power in the same frequency band. To trace the temporal positions of the animals, two color LEDs were set on the headstage and were recorded by digital video camera at 30 frames/s. Spikes were extracted from the high-pass filtered signals (median filter, cut-off frequency: 800 Hz). Spike sorting was performed semi-automatically, using KlustaKwik2 (Kadir et al., 2014). The cell-types of the units were classified by peak-trough latency and width. In total, we analyzed 831 putative excitatory neurons (n = 639 for wildtype; n = 123 for cKO; n = 69 for NR1f/f mice Table S1 and S4) and 250 inhibitory neurons (n = 169 for wildtype; n = 53 for cKO; n = 28 for NR1f/f mice). The positions of the animals were determined by the position of the LEDs mounted on the headstage by custom-made tracking software. The rate maps of the spike number and occupancy probability was generated from 4-cm binned segments from the position and spiking data. The normalized PSTH for individual neurons in delay-act and delay–sup cells in the CA1 was computed under delay 20 s conditions. The autocorrelation of the population vector was then computed.

### Determination of delay-active and delay-suppressive neurons

To examine the effect of delay on neuronal activities, we quantified changes of the firing rate of each neuron during the long delay period. First, we calculated the firing rate in the delay zone (*R_delay_* = spike number in delay zone *n_delay_* / time spent in delay zone *t_delay_;* see Figure 1A), and that in all zones (*R_total_* = spike number in all zones *n_total_* / time spent in all zones *t_total_*) in long-delay trials, and then computed the ratio of them (*R_delay_* / *R_total_*). Second, we performed a permutation test in order to determine whether the ratio of the firing rates *R_delay_* / *R_total_* shows significant change or not. To make surrogate data, we resampled the spike trains by permuting the inter-spike-intervals and by realigning with them. We repeated this process 1000 times to obtain 1000 resampled datasets. The rank of the original firing rate ratio *R_delay_*/ *R_total_* in the resampled 1000 firing rate ratios was used to define the statistical assessment (delay-act cells: significant higher firing rate (rank < 50, top 5 %); delay-sup cells: significant lower firing rate (rank > 950, bottom 5 %)).

### Statistical analysis

Correlation coefficients and p values between firing rates and delay length were calculated by the Matlab function. To compare the firing rates between short and long delay conditions, we performed Wilcoxon’s rank sum test. To assess side-dependency in firing rates, three-way ANOVA [side (right and left) × phase (start, delay, and goal) × delay length (5 and 20)] was used. To compare the effect of devaluation or revaluation on firing rate of delay-act and delay-sup cells, Mann-Whitney U test was carried out. To examine the ratio distribution, we performed two-tailed one-sample t tests against 0. The behavioral impact of NMDAR conditional knockout was evaluated by two-way ANOVA [genotype (cKO and control) × choice probability] followed by post hoc Scheffe’s test. Fisher’s exact test was applied to compare the cell-type distributions between cKO and control mice.

## Acknowledgments

We thank Drs. Steven Middleton, Roman Boehringer, and Chinnakkaruppan Adaikkan for help building tetrodes, Dr. Charles Yokoyama for valuable comments, and Dr. Susumu Tonegawa for providing us NR1 cKO mice. Funding was provided by Grand-in-Aid for Exploratory Research (JSPS KAKENHI Grant Number 16K15196) and the “Brain/MINDS” from Japan Agency for Medical Research and Development (AMED).

## Author Contributions

A.M., S.F., and S.I. designed the experiments. Q.Z., H.G., and S.I. developed the automated T-maze. A.M. and C.S. performed experiments. A.M. and S.F. performed data analysis. A.M., S.F., T.M., and S.I. prepared the paper.

## Supplemental figure titles and legends

Figure S1 related to Figure 2-8. Histological verification of recording sites in the CA1.

(A) An example of hippocampal slice with DiI pasted at tip of the electrode and DAPI staining. White arrows indicate inserted points.

(B) Electrode tracks are represented by black dots.

Figure S2 related to Figure 2-8. Cell-type classification by plot in peak-trough and firing rate 2D dimension. Red triangles indicate excitatory neurons, while blue circles indicate inhibitory neurons. Dotted line indicates classification criteria for excitatory and inhibitory determinations.

Figure S3 related to Figure 3-5. Temporal patterns of the CA1 neuron shift as sessions progressed.

(A) Firing gradually shifted backward.

(B) Comparison of firing patterns between pre– and post-long delay-session(s).

(C) Normalized firing rate sorted by peak firing rate time on pre-(Left) and post-long-delay-session (Center). The order of the cells in the pre– and post was identical. Difference in peak firing time between pre– and post-long-delay-session (sorted by time, Right).

Figure S4 related to Figure 7. Firing in CA1 delay-act cells depended on animal preference. Forced conditions where the animals were forced to choose one side by a partition (here, left side). Yellow line indicates the trial converted to forced conditions.

(A) The firing in the left side disappeared when the preference to the left side was extinguished.

(B) Simultaneously, other neurons showed increased firing when the preference recovered to the other side (here, right side).

Figure S5 related to Figure 8. Mutant mice (CA1-NR1cKO) exhibiting impaired delay discounting showed less specific spatial representation in place cell activities and less in the number of delay-active cells.

(A) Spatial representation of example neurons from delay-act in CA1-NR1cKO (upper) and control mice (lower). Arrows indicate areas with non-specific firing.

(B) Percentages of delay-act and -sup cells in mutant and control mice. Error bars indicate 95% Clopper-Pearson’s confidence intervals.

Table S1 related to Figure 3-7. The number of delay-active and delay-suppressed CA1 excitatory and inhibitory neurons recorded from all sessions.

Table S2 related to Figure 3-7. Full distribution of CA1 excitatory neurons for all the tested conditions.

Table S3 related to Figure 6. Distribution of side-dependent and side–independent, delay-active and delay-suppressive, excitatory and inhibitory neurons.

Table S4 related to Figure 8. Full distribution of CA1 excitatory neurons for the NMDAR mutant study. The numbers with parentheses are cells from the wildtype.

