## Supplementary figures and images for "The hippocampus encodes delay and value information during delay-discounting decision making"

### Figure S1

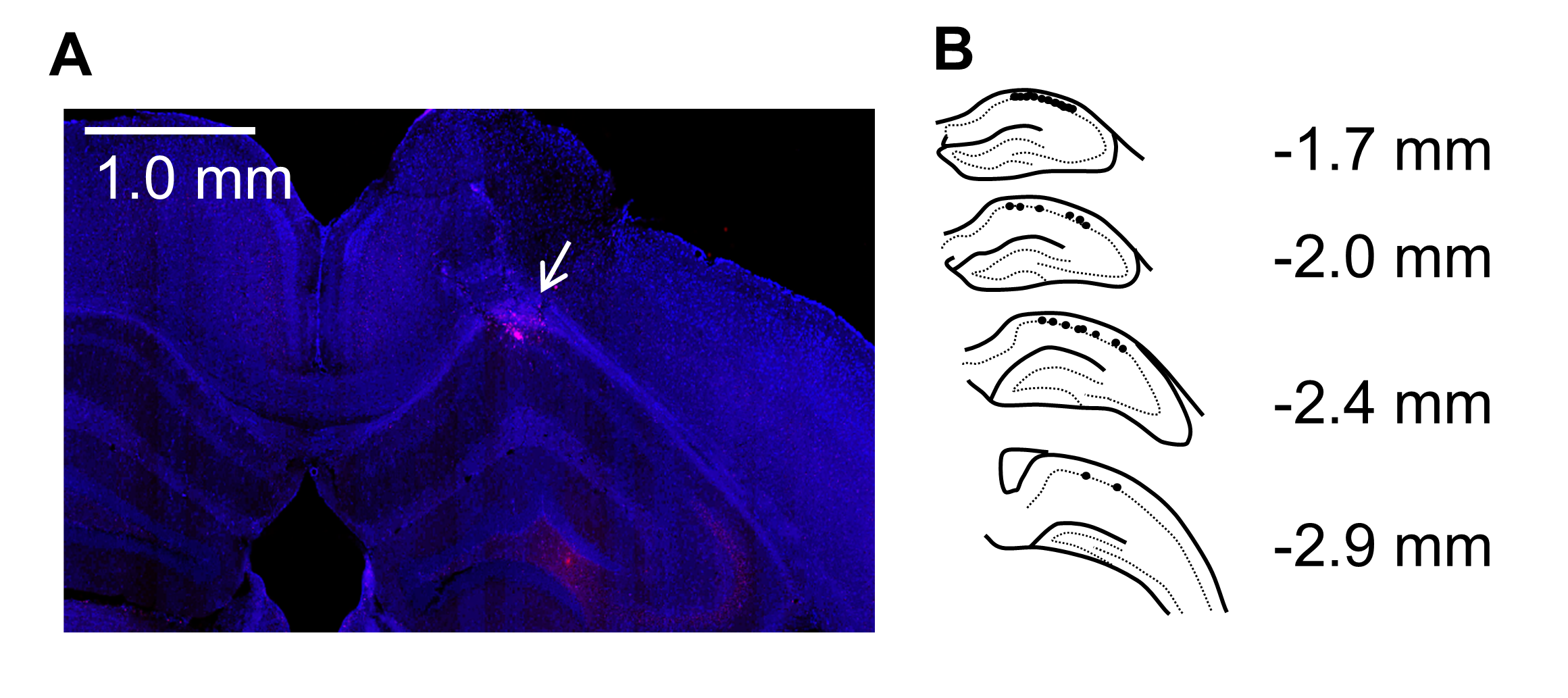

### Figure S2

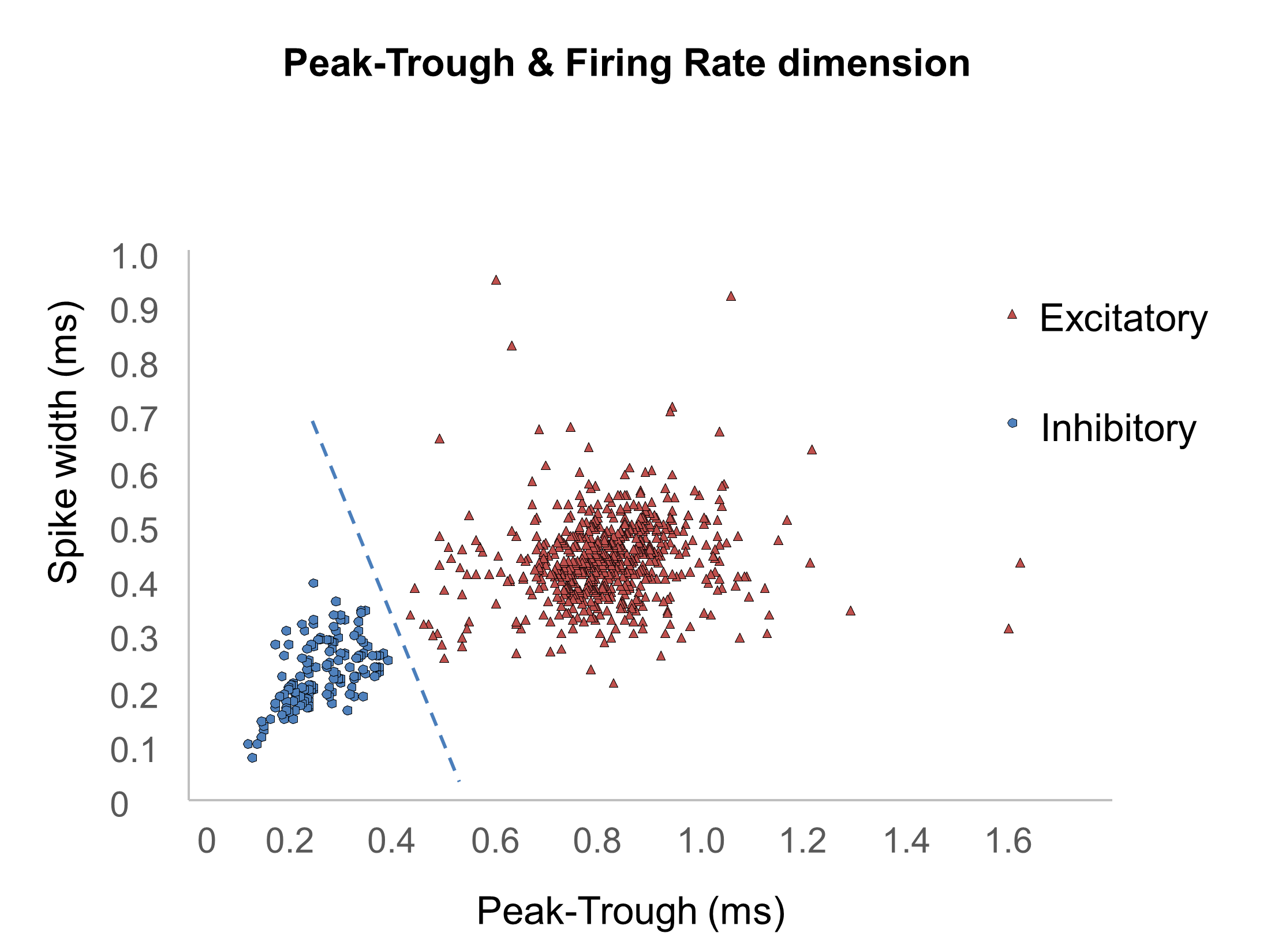

### Figure S3

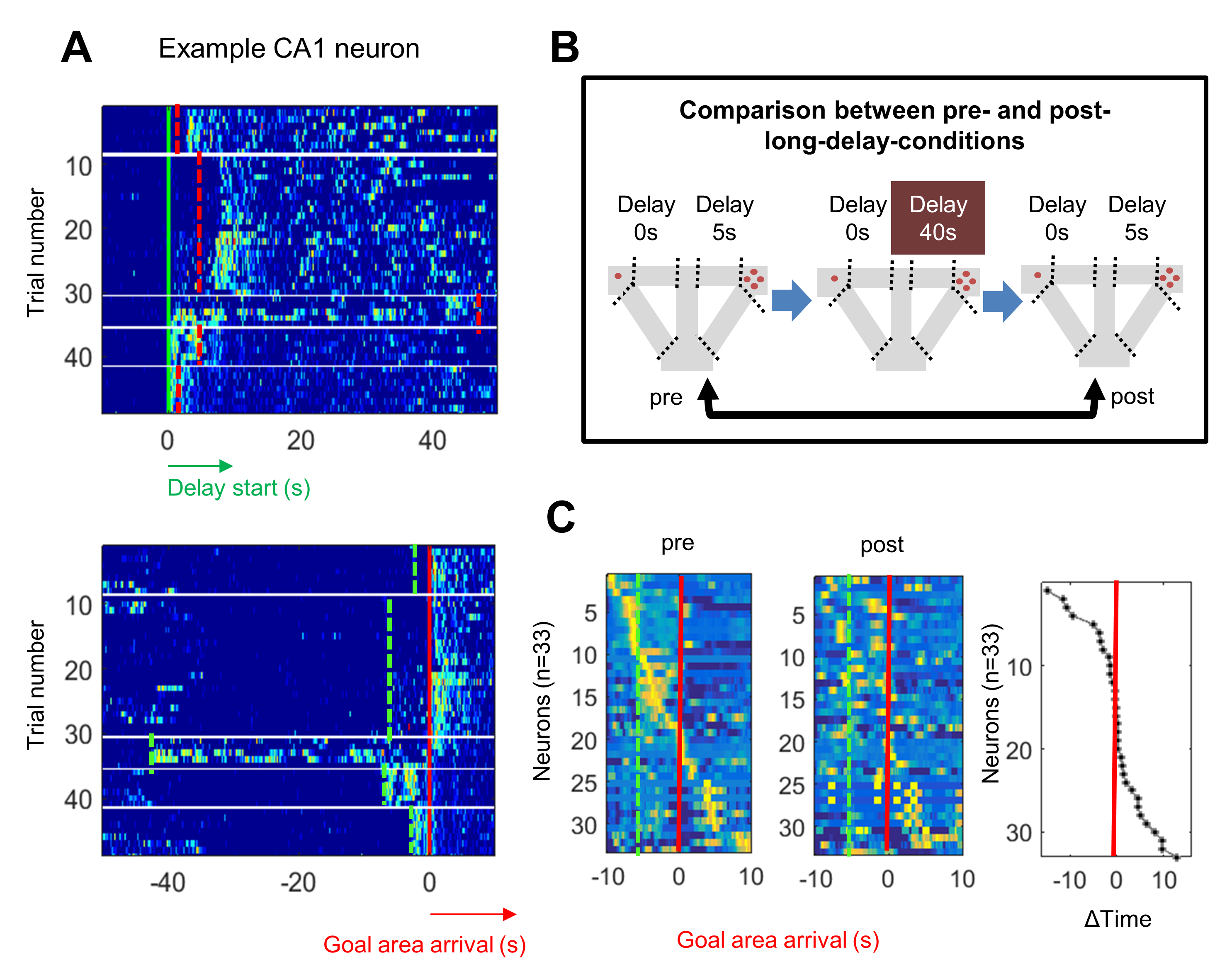

### Figure S4

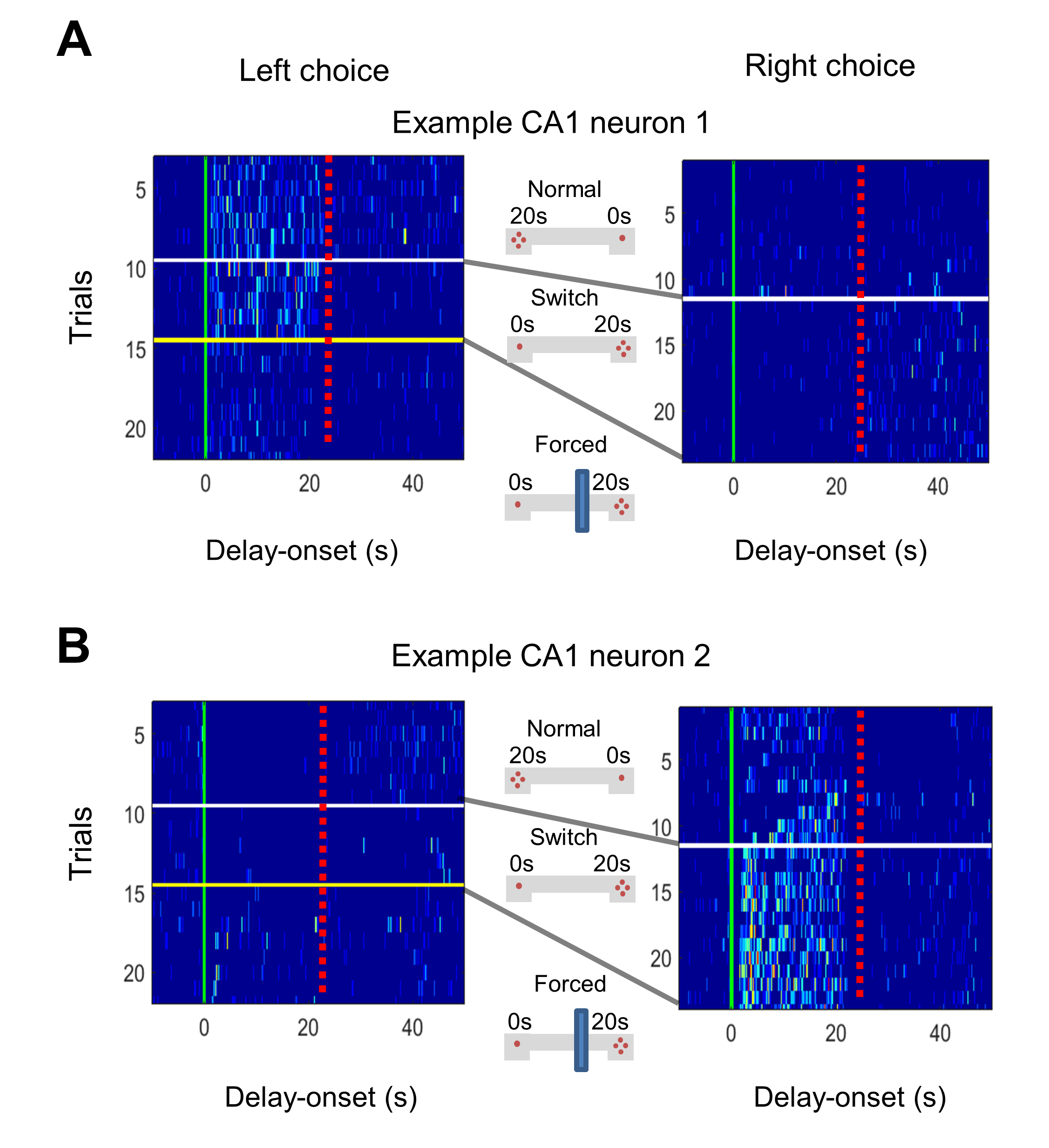

### Figure S5

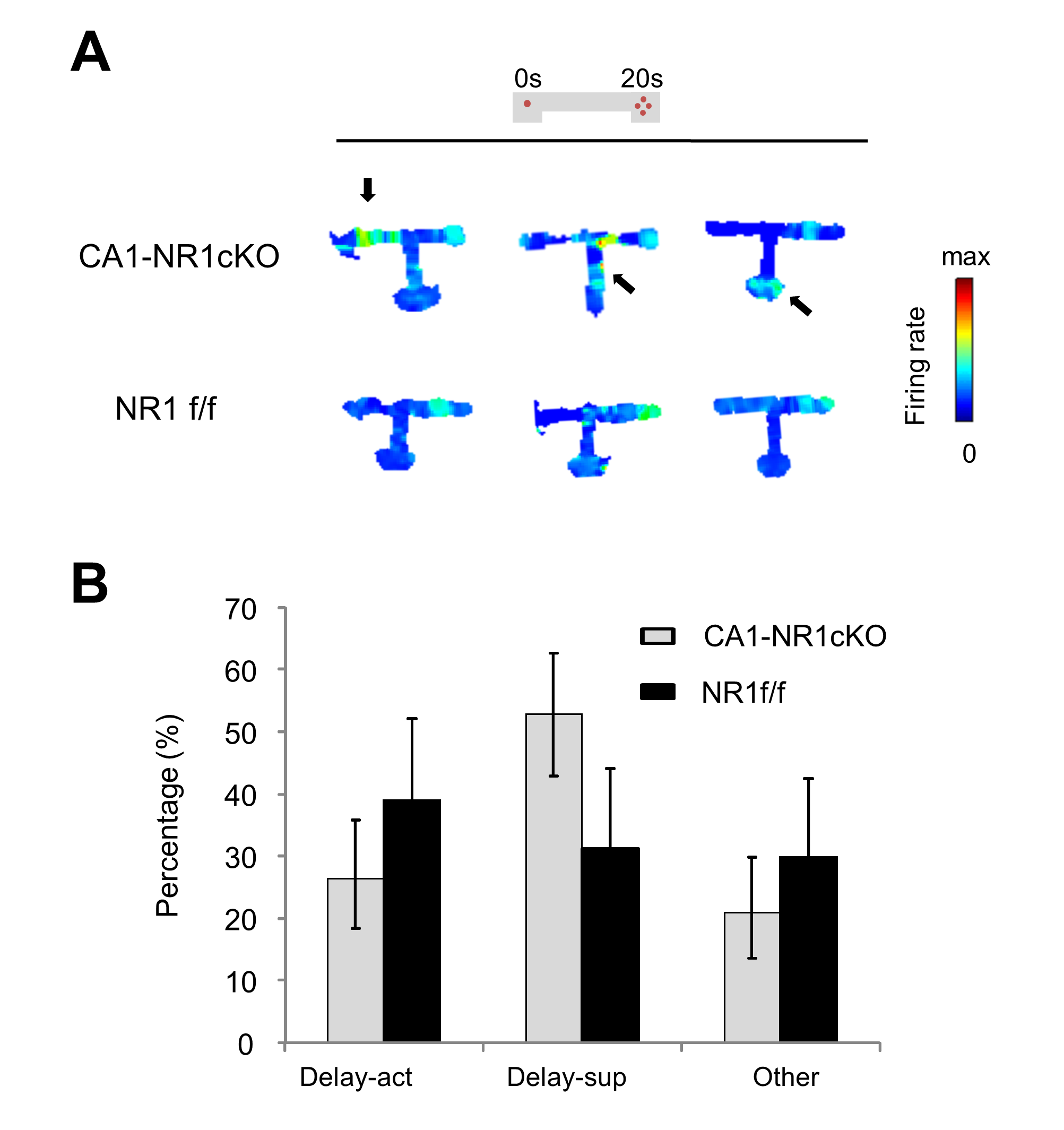

### Table S1

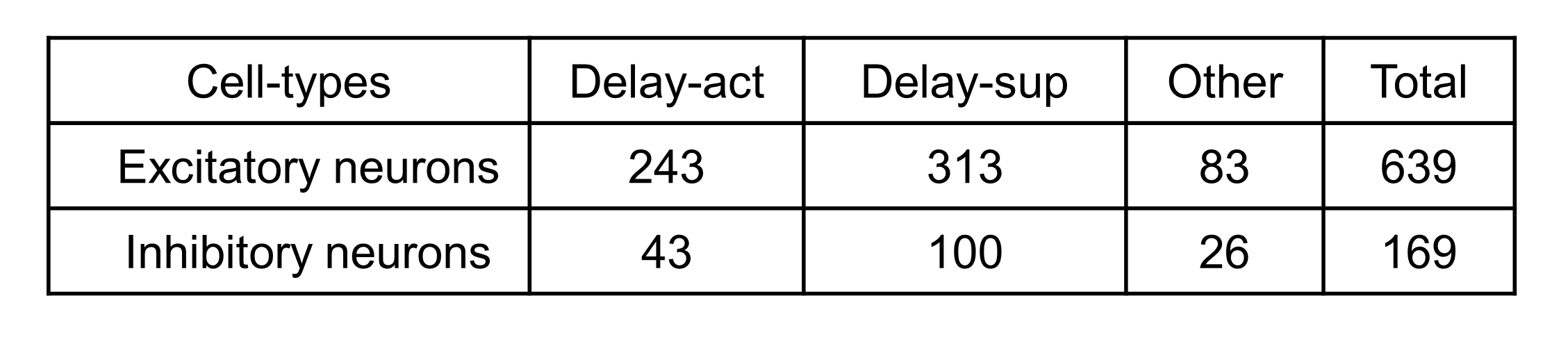

### Table S2

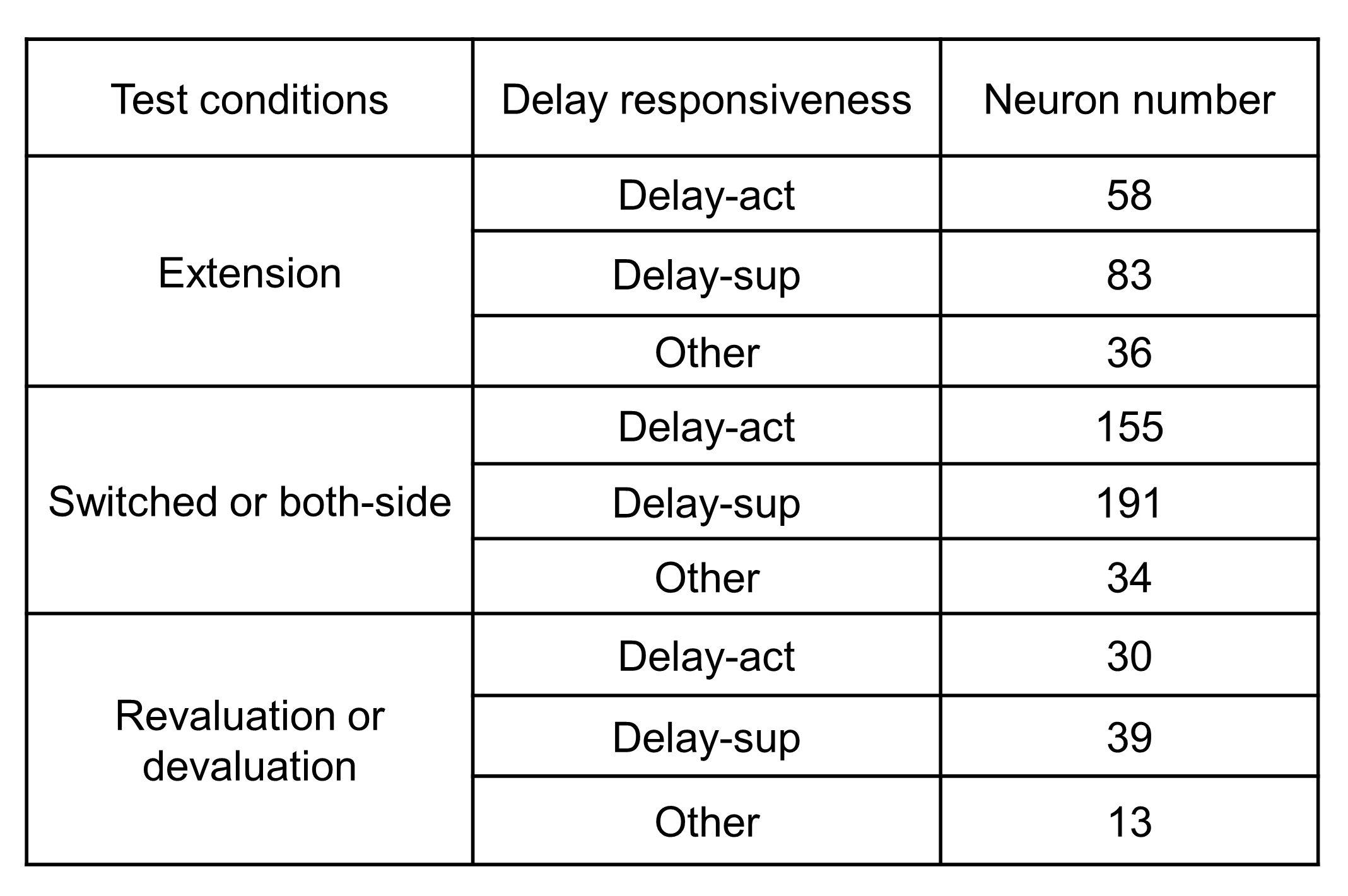

### Table S3

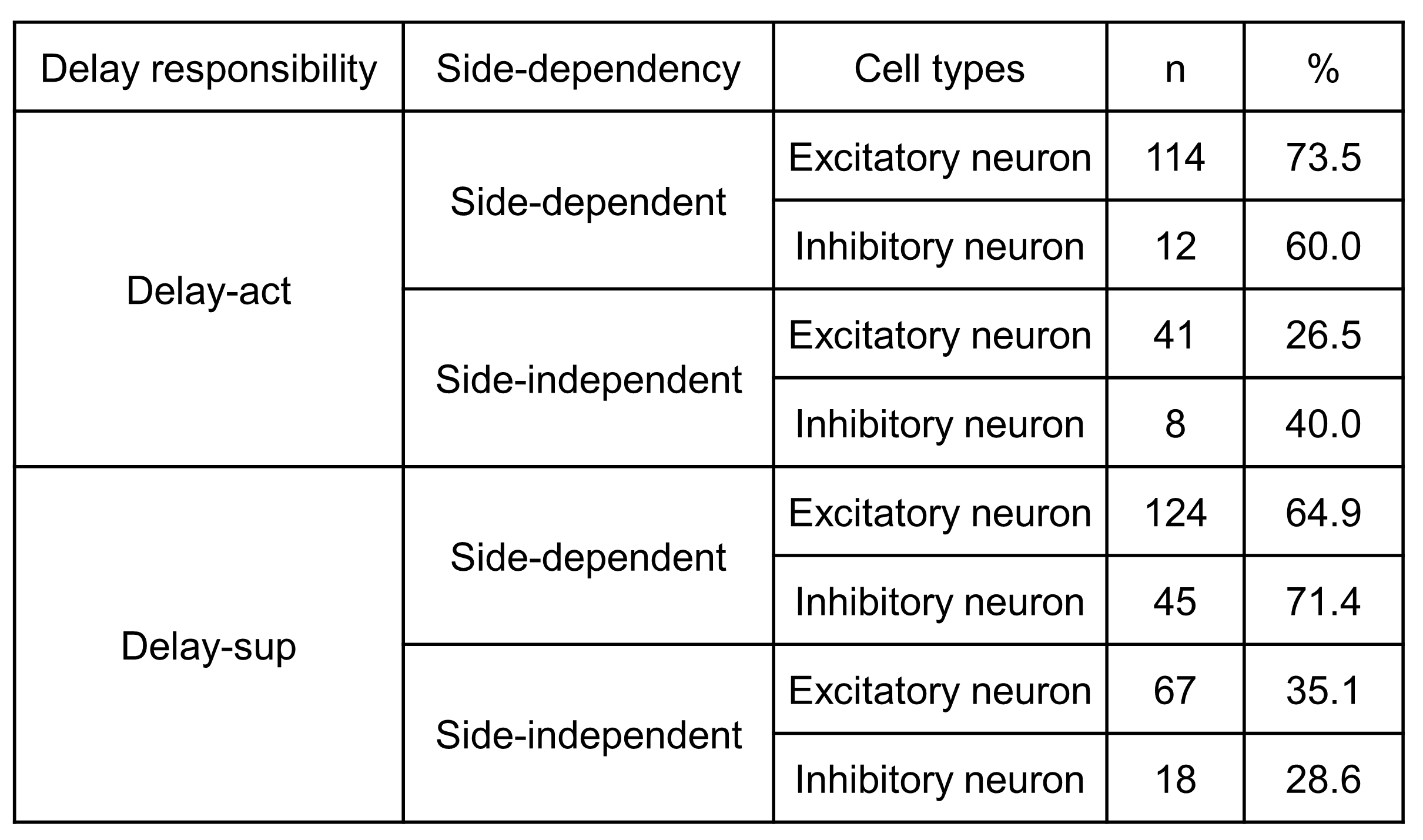

### Table S4

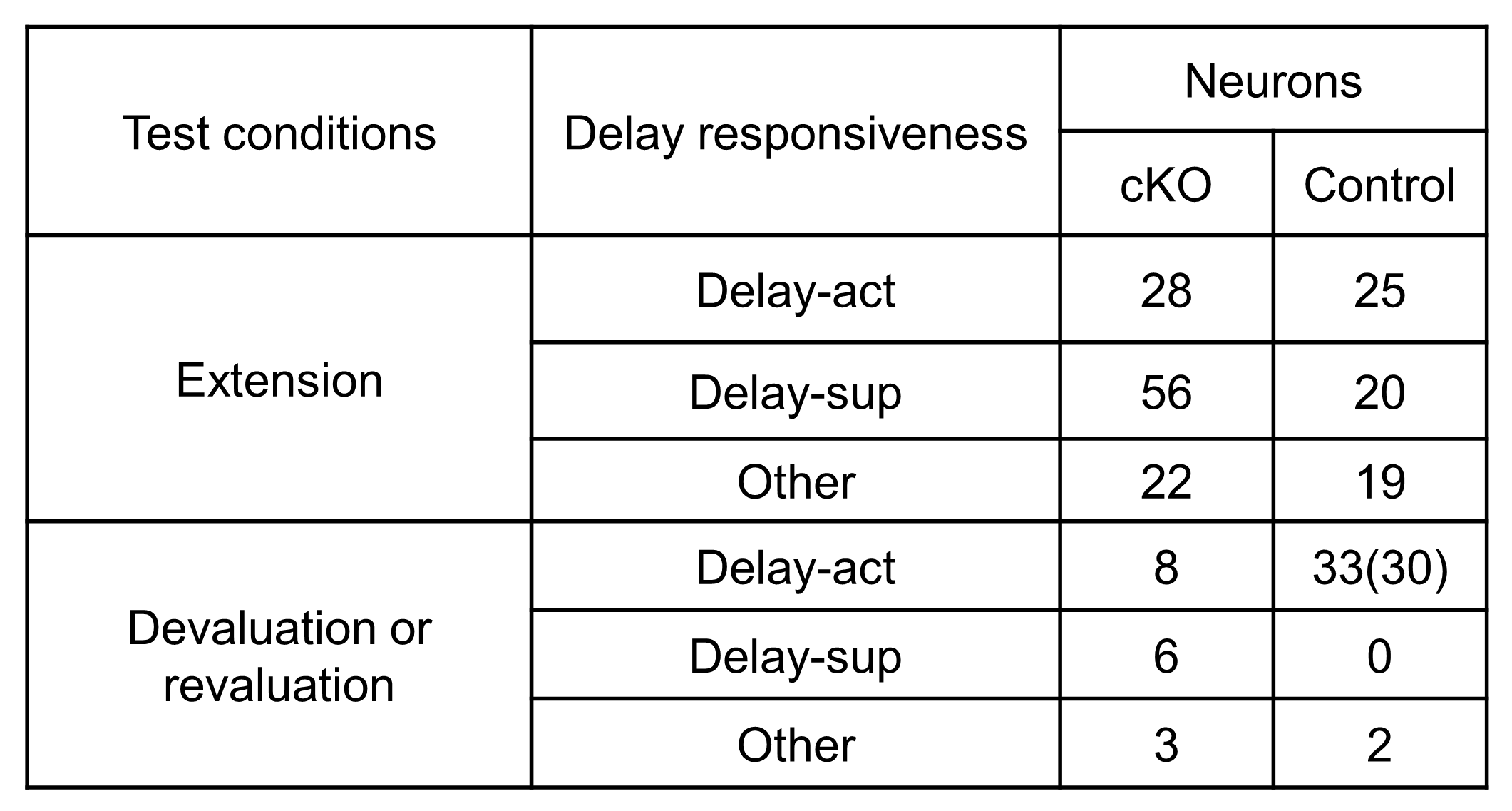
